# Neutralization of SARS-CoV-2 Omicron BA.2 by Therapeutic Monoclonal Antibodies

**DOI:** 10.1101/2022.02.15.480166

**Authors:** Hao Zhou, Takuya Tada, Belinda M. Dcosta, Nathaniel R. Landau

## Abstract

Monoclonal antibody therapy for the treatment of SARS-CoV-2 infection has been highly successful in decreasing disease severity; however, the recent emergence of the heavily mutated Omicron variant has posed a challenge to this treatment strategy. The Omicron variant BA.1 has been found to evade neutralization by several of the therapeutic monoclonal antibodies authorized for emergency use, while Vir-7831 and a cocktail consisting of monoclonal antibodies AZD8895+AZD1061 retain significant neutralizing activity. A newly emerged variant, Omicron BA.2, containing some of the BA.1 mutations plus an additional 6 mutations and 3 deletions, 3 of which lie in the receptor binding domain, has been found to be spreading with increased transmissibility. We report here, using spike protein-pseudotyped lentiviruses, decreased neutralization of BA.2 by several therapeutic monoclonal antibodies but that the mixture of AZD8895+AZD1061 retained substantial neutralizing activity against BA.2.

## Main text

The BA.2 variant of the highly mutated Omicron SARS-CoV-2 variant was identified in late 2021 in patients in countries including Denmark, South Africa and India (1) and since then, in more countries including the United States. BA.2 has been found to be 1.5-fold more transmissible than the already highly transmissible BA.1 variant and is thus expected to continue to increase in prevalence (2). The BA.2 spike protein has all of the mutations of BA.1 plus an additional 6 mutations and 3 deletions, three of which lie in the receptor binding domain (3) (**Supplemental Figure 1A**). Of the monoclonal antibodies authorized by the Food and Drug Administration (FDA) for emergency use (4), Regeneron REGN10933 and REGN10987 and Eli Lilly LY-CoV555 and LY-CoV016 were found to be largely inactive against BA.1 while the GlaxoSmithKline/Vir monoclonal antibody Vir-7831 (5) and AstraZeneca Evusheld cocktail consisting of monoclonal antibodies AZD8895 and AZD1061 retain considerable neutralizing titer BA.1 (6–8). The Evusheld cocktail of AZD8895+AZD1061 is formulated for slow release for use prophylactically primarily in immunocompromised individuals (9).

**Figure 1.**
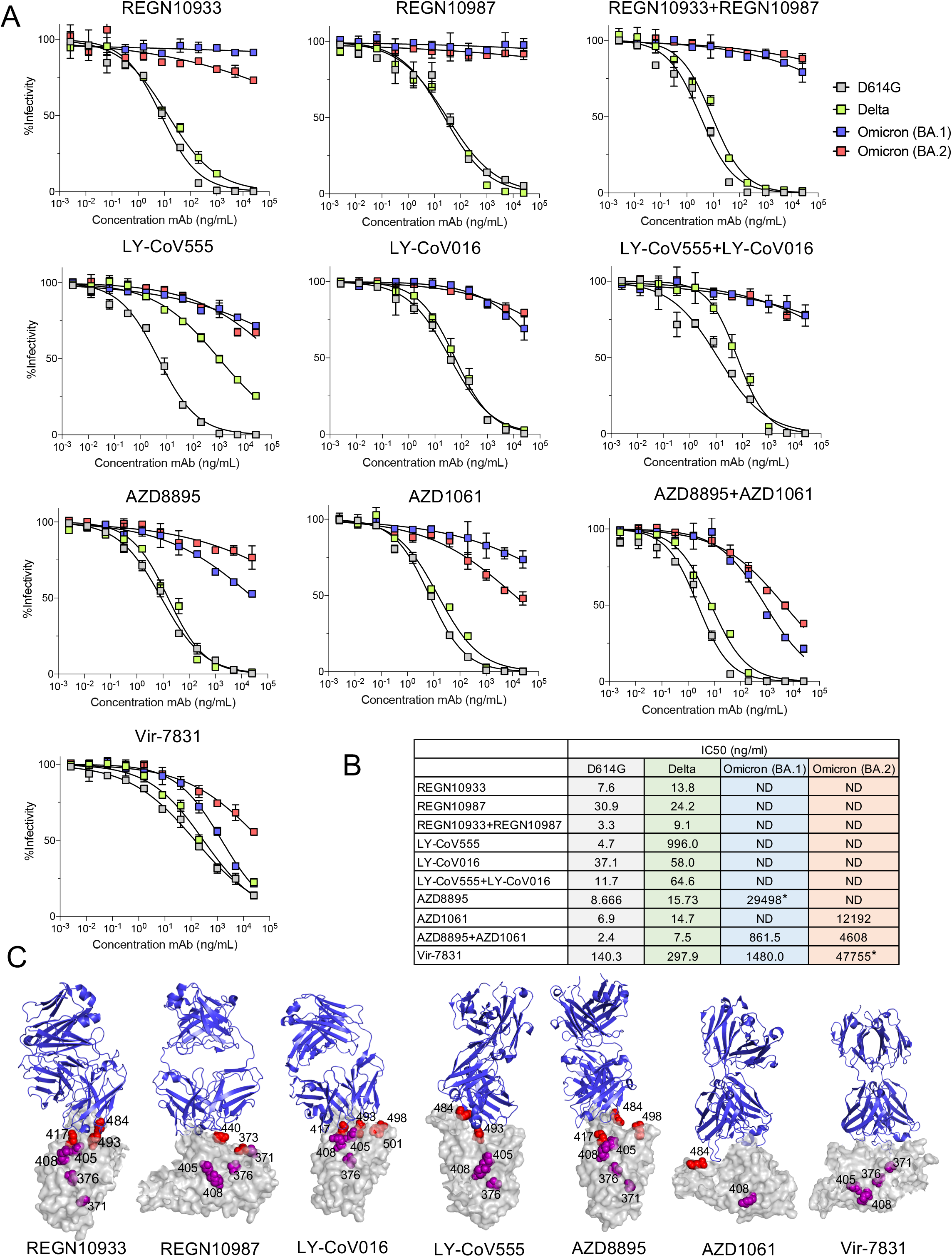
Neutralization of BA.2 by therapeutic monoclonal antibodies. (A) Lentiviral pseudotyped viruses were generated as previously described using codon-optimized 19 amino acid deleted spike proteins (15). A fixed amount of virus, normalized for reverse transcriptase activity, was treated with 5-fold serially diluted monoclonal antibody, in duplicate, for 30 minutes and then used to infect ACE2.293T cells. Luciferase activity was measured after 24 hours and the data are plotted as curves neutralized to luciferase activity in the absence of antibody. (B) The IC_50_ was calculated from the neutralization curves by GraphPad Prism 8 software. Values that approached 50% neutrlization were estimated, as indicated by an asterisk (*); those that did not approach 50% neutralization were not determined (ND). (C) The location of mutations found to affect monoclonal antibody binding is shown on antibody: spike protein complexes. Complexes are visualized with PyMOL Molecular Graphics System, v2.1.1 (Schrödinger, LLC) software. Mutations in BA.1 (that are also in BA.2) previously reported to have >5-fold effect on neutralizing titer are shown in red. Mutations identified in this study with >2-fold effect are shown in purple.

In this report, we tested the ability of the therapeutic monoclonal antibodies to neutralize the BA.2 variant, with particular interest in Vir-7831 and AZD8895+AZD1061 given their ability to neutralize BA.1. Neutralizing antibody titers against the parental D614G variant, Delta, BA.1 and BA.2 were determined using spike protein-pseudotyped lentiviruses. The lentiviral pseudotype assay has been found to yield neutralizing antibody titers that are in close agreement with those determined with the live virus plaque reduction neutralization test (PRNT) (10). To understand the mechanism by which the variants evade monoclonal antibody neutralization, we tested viruses with spike proteins containing the individual RBD point mutations of BA.2 for their effect on neutralization by each antibody.

Neutralization curves and the IC_50_s calculated from the curves for D614G, Delta, BA.1 and BA.2 pseudotyped viruses are shown in **Figures 1A and B**. The monoclonal antibodies potently neutralized the D614G virus and neutralized the Delta variant except for LY-CoV555 (Bamlanivimab) for which the neutralizing titer was decreased 39-fold compared to D614G, consistent with previous reports (11). REGN10933, REGN10987, REGN10933+REGN10987, LY-CoV555, LY-CoV016, LY-CoV555 + LY-CoV016, AZD8895, AZD1061 lacked detectable neutralizing titer against BA.1. The AZD8895+AZD1061 mixture and the Vir-7831 monoclonal antibody neutralizing titers were decreased by 359-fold and 11-fold compared to D614G, consistent with previous reports (6–8). REGN10933, REGN10987, REGN10933+REGN10987, LY-CoV555, LY-CoV016, and LY-CoV555+LY-CoV016 lacked detectable neutralizing titer against BA.2 while Vir-7831 was active against BA.2, albeit with decreased titer. In contrast, AZD1061 retained considerable neutralizing titer against BA.2 and AZD8895, which had little activity on its own, synergized with AZD1061 to restore much of the neutralizing titer against BA.2 in the AZD8895+AZD1061 mixture (**Figure 1A and B**).

Analysis of viruses pseudotyped by spike proteins with the individual novel RBD mutations of BA.2 provided insight into the cause of the decrease in neutralization titers by the monoclonal antibodies (**Supplemental Figure 1B and C**). The structural models of the spike protein: antibody complexes are shown with mutations previously found to cause >5-fold decrease in titer against mutations present in both BA.1 and BA.2 as well as novel the BA.2 mutations identified here that cause >2-fold decreased in titer. For the most part, the active mutations in BA.1 map to the sites of interaction with the antibodies while the additional active mutations in BA.2 (S371F, T376A, D405N and R408S) lie distal to the interaction sites (**Figure 1C**). Presumably, these mutations act at a distance by affecting the conformation of the RBD spike.

## Discussion

We report here that while several therapeutic monoclonal antibodies have lost neutralizing titer against both BA.1 and BA.2, the combination of AZD8895+AZD1061 retained considerable neutralizing titer against both Omicron subtypes in a lentiviral pseudotype assay (6–8). The increased neutralizing titer of the mixture against BA.2 appeared to result from synergy between the two monoclonal antibodies.

Since the identification of the Omicron BA.2 variant in several countries including Denmark, South Africa and India, the variant has been identified in infected individuals in additional countries and mathematical models predict that its increased transmissibility will result in its continued increase in prevalence (2). While monoclonal antibody therapy has been highly effective at preventing hospitalization and death, the emergence of the Omicron variant poses a major threat to the efficacy of current treatments. As BA.2 prevalence increases, current monoclonal antibodies may become less effective for the treatment of COVID-19. Therapies that target BA.2 and potential future variants that may emerge are therefore of great importance. The findings presented here demonstrate the difficulty of finding a pan-neutralizing monoclonal antibody against SARS-CoV-2. Our findings support the importance of small molecule drugs such as Molnupiravir (12) and Nirmatrelvir (13) that act on viral targets outside of the highly mutable spike protein (14). They also provide further rationale to protect against severe disease by vaccination which induces cross-reactive antibodies and T cell responses against spike protein epitopes that are not mutated in the Omicron variants.

## Material and Methods

### Plasmids

The pCMV3-SARS-CoV-2-BA.2-Spike was kindly provided by David Ho’s group. Spike expression vectors with the individual mutations of the Omicron BA.2 RBD were generated by overlap PCR mutagenesis using the D614G spike expression vector pcCOV2.∆19.D614G as a template.

### Cells

293T and ACE2.293T were grown in Dulbecco’s Modified Eagle’s Medium with 10% fetal bovine serum at 37°C and 5% CO_2_.

### SARS-CoV-2 spike proteins lentiviral pseudotypes

SARS-CoV-2 spike protein pseudotyped lentivirus stocks were produced by cotransfection of 293T cells with pMDL Gag/Pol vector, plenti.GFP.nLuc and spike protein expression vectors which deleted 19 amino acid from cytoplasmic tail (15). After 2 days, supernatants were harvested and concentrated by ultracentrifugation. The viruses were normalized for reverse transcriptase (RT) activity.

### Antibody neutralization assay

Monoclonal antibodies were serially diluted (5-fold) and incubated with pseudotypes (MOI=0.2 on target cells). After 30 minutes incubation, the virus was added to target cells in a 96 well culture dish. After 2 days of infection, infectivity was developed by Nano Glo substrate and luminescence was read in an Envision 2103 microplate luminometer.

### Data analysis

All samples were tested in duplicate. Data were analyzed using GraphPad Prism 8 software. Analyses of the structures of the SARS-CoV-2 spike protein with antibody Fabs was performed with the PyMOL Molecular Graphics System, v2.1.1 (Schrödinger, LLC). The PDB accession codes for the structures shown are 6XDG (REGN10933 and REGN10987), 7KMG (LY-CoV555), 7C01 (LY-CoV016), 7R6W (Vir-7831), and 7L7E (AZD8895 and AZD1061).

## Acknowledgements

We thank Sho Iketani and David D. Ho (Columbia University Vegelos College of Physicians and Surgeons) for the BA.2 expression vector. The work was funded by grants from the NIH to N.R.L. (DA046100, AI122390 and AI120898).

---

**Supplementary Figure 1.**
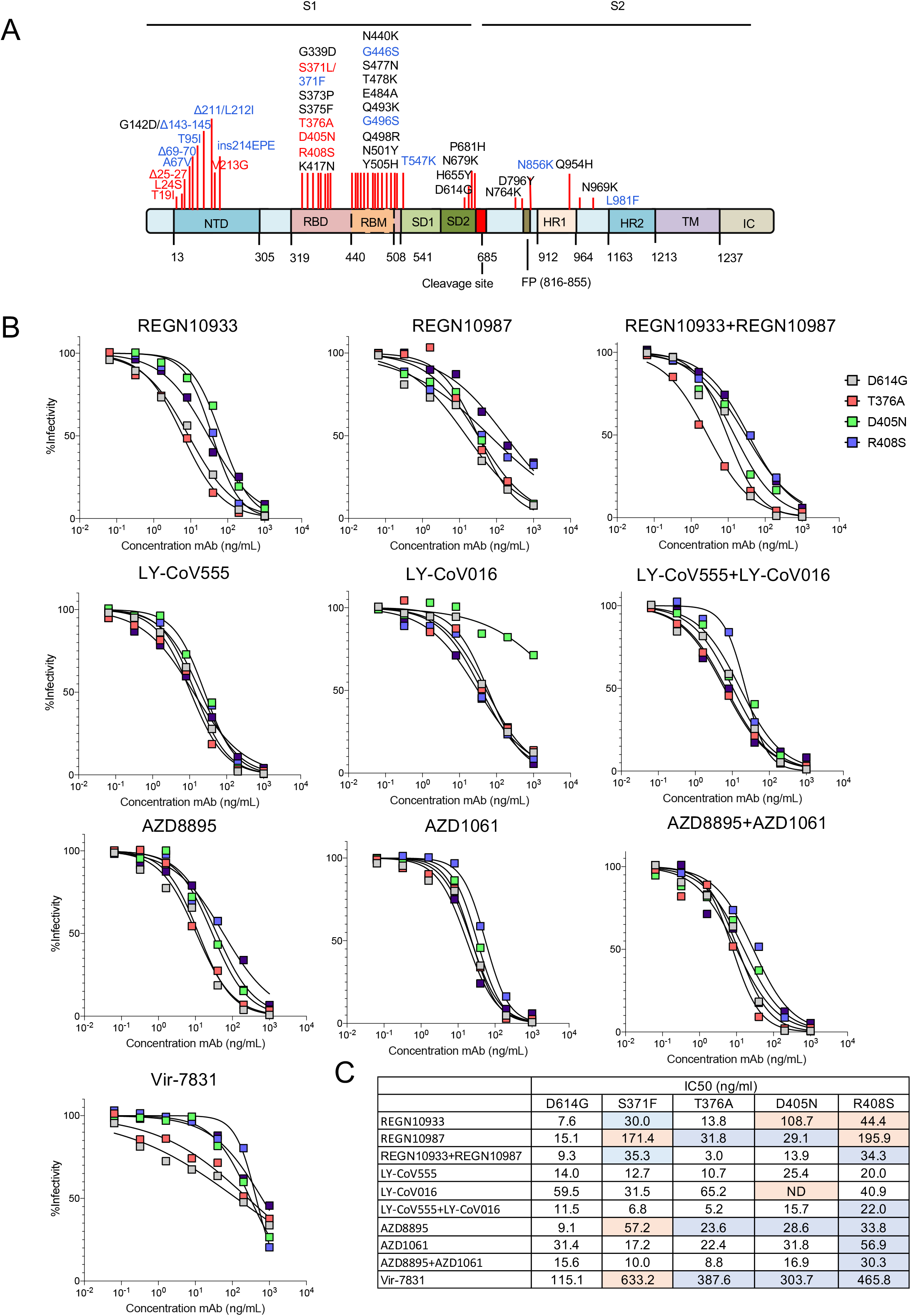
Neutralization of single point mutated virus (BA.2) by therapeutic monoclonal antibodies. (A) The structure of the SARS-CoV-2 Omicron BA.2 spike is indicated. NTD, N-terminal domain; RBD, receptor-binding domain; RBM, receptor-binding motif; SD1 subdomain 1; SD2, subdomain 2; CS, cleavage site; FP, fusion peptide; HR1, heptad repeat 1; HR2, heptad repeat 2; TM, transmembrane region; IC, intracellular domain. Novel mutations found in BA.2 are shown in red. The mutations which are specific to BA.1 are shown in blue. (B) Individual point mutated viruses were treated with 5-fold serially diluted monoclonal antibodies and then used to infect ACE2.293T cells. Luciferase activity was measured after 24 hours. (C) The IC_50_ was calculated from the neutralization curves using GraphPad Prism 8 software. The point mutations found to cause >5-fold decrease in neutralizing titer are shown in red. The point mutations found to cause >2-fold decrease in neutralizing titer are shown in blue. Values that did not approach 50% neutralization were not determined (ND).

## References

1. CDC. Omicron Variant: What You Need to Know. 2022.

2. Lyngse FP, Kirkeby CT, Denwood M, Christiansen LE, Mølbak K, Møller CH, et al. Transmission of SARS-CoV-2 Omicron VOC subvariants BA.1 and BA.2: Evidence from Danish Households. medRxiv. 2022:2022.01.28.22270044.

3. Majumdar S, Sarkar R. Mutational and phylogenetic analyses of the two lineages of the Omicron variant. J Med Virol. 2021 Dec 29.

4. FDA. 2022: https://www.fda.gov/news-events/press-announcements/coronavirus-covid-19-update-fda-limits-use-certain-monoclonal-antibodies-treat-covid--due-omicron.

5. Group AC-TfIwC-S. Efficacy and safety of two neutralising monoclonal antibody therapies, sotrovimab and BRII-196 plus BRII-198, for adults hospitalised with COVID-19 (TICO): a randomised controlled trial. Lancet Infect Dis. 2021 Dec 23.

6. Tada T, Zhou H, Dcosta BM, Samanovic MI, Chivukula V, Herati RS, et al. Increased resistance of SARS-CoV-2 Omicron Variant to Neutralization by Vaccine-Elicited and Therapeutic Antibodies. bioRxiv. 2021:2021.12.28.474369.

7. VanBlargan L, Errico J, Halfmann P, Zost S, Crowe J, Purcell L, et al. An infectious SARS-CoV-2 B.1.1.529 Omicron virus escapes neutralization by therapeutic monoclonal antibodies. Res Sq. 2021 Dec 27.

8. Liu L, Iketani S, Guo Y, Chan JFW, Wang M, Liu L, et al. Striking Antibody Evasion Manifested by the Omicron Variant of SARS-CoV-2. Nature. 2021 2021/12/23.

9. ClinicalTrials.gov. phase III Double-blind, Placebo-controlled Study of AZD7442 for Pre-exposure Prophylaxis of COVID-19 in Adult. (PROVENT).

10. Noval MG, Kaczmarek ME, Koide A, Rodriguez-Rodriguez BA, Louie P, Tada T, et al. Antibody isotype diversity against SARS-CoV-2 is associated with differential serum neutralization capacities. Scientific Reports. 2021 2021/03/10;11(1):5538.

11. Planas D, Veyer D, Baidaliuk A, Staropoli I, Guivel-Benhassine F, Rajah MM, et al. Reduced sensitivity of SARS-CoV-2 variant Delta to antibody neutralization. Nature. 2021 2021/08/01;596(7871):276–80.

12. Jayk Bernal A, Gomes da Silva MM, Musungaie DB, Kovalchuk E, Gonzalez A, Delos Reyes V, et al. Molnupiravir for Oral Treatment of Covid-19 in Nonhospitalized Patients. N Engl J Med. 2022 Feb 10;386(6):509–20.

13. Owen DR, Allerton CMN, Anderson AS, Aschenbrenner L, Avery M, Berritt S, et al. An oral SARS-CoV-2 M^pro^ inhibitor clinical candidate for the treatment of COVID-19. Science. 2021;374(6575):1586–93.

14. Li P, Wang Y, Lavrijsen M, Lamers MM, de Vries AC, Rottier RJ, et al. SARS-CoV-2 Omicron variant is highly sensitive to molnupiravir, nirmatrelvir, and the combination. Cell Research. 2022 2022/01/20.

15. Tada T, Fan C, Chen JS, Kaur R, Stapleford KA, Gristick H, et al. An ACE2 Microbody Containing a Single Immunoglobulin Fc Domain Is a Potent Inhibitor of SARS-CoV-2. Cell Rep. 2020 Dec 22;33(12):108528.

